# Transcriptome stability profiling using 5’-bromouridine IP chase (BRIC-seq) identifies novel and functional microRNA targets in human melanoma cells

**DOI:** 10.1101/555540

**Authors:** Piyush Joshi, Tatsuya Seki, Shinobu Kitamura, Andrea Bergano, Bongyong Lee, Ranjan J. Perera

**Affiliations:** Department of Oncology, Johns Hopkins University School of Medicine, 401 N. Broadway, Baltimore, MD 21287 USA; Cancer and Blood Disorders Institute, Johns Hopkins All Children’s Hospital, 600 5th St. South, St. Petersburg, FL 33701 USA; Sanford Burnham Prebys Medical Discovery Institute, 6400 Sanger road, Orlando, FL 32827 USA; Medical and Biological Laboratories, Nagoya 460-0008, Japan; The Sidney Kimmel Comprehensive Cancer Center, Johns Hopkins University School of Medicine, 401 N. Broadway, Baltimore, MD 21287 USA

**Keywords:** BRIC-seq, RNA-seq, melanoma, microRNA, RNA stability

## Abstract

RNA half-life is closely related to its cellular physiological function, so stability determinants may have regulatory functions. Micro(mi)RNAs have primarily been studied with respect to post-transcriptional mRNA regulation and target degradation. Here we study the impact of the tumor suppressive melanoma miRNA miR-211 on transcriptome stability and phenotype in the non-pigmented melanoma cell line, A375. Using 5’-bromouridine IP chase (BRIC)-seq, transcriptome-wide RNA stability profiles revealed highly regulated genes and pathways important in this melanoma cell line. By combining BRIC-seq, RNA-seq and *in silico* predictions, we identified both existing and novel direct miR-211 targets. We validated *DUSP3* as one such novel miR-211 target, which itself sustains colony formation and invasion in A375 cells via MAPK/PI3K signaling. miRNAs have the capacity to control RNA turnover as a gene expression mechanism, and RNA stability profiling is an excellent tool for interrogating functionally relevant gene regulatory pathways and miRNA targets when combined with other high-throughput and *in silico* approaches.

## INTRODUCTION

Melanoma is the commonest cause of skin cancer-related deaths, with an estimated 90,000 diagnoses and 10,000 deaths in the US in 2018 [1]. The high mortality associated with melanoma is due to its intrinsic aggressiveness, metastatic potential, and resistance to conventional therapies [2]. There is a pressing need to better understand the abnormal biological processes contributing to melanoma development, metastasis, and resistance to develop new biomarkers and therapies.

Whole transcriptome sequencing (RNA-seq) is now a standard method to interpret the cellular state during disease development and progression. However, RNA-seq represents a static image of the process of transcription, while biological processes are by their very nature dynamic. Recent studies on transcriptome dynamics have shown that transcript stability is correlated with function [3–6]. For instance, transcription factor mRNAs have high average decay rates [6] and mRNA turnover correlates with signaling pathways active in cancer, suggesting a post-transcriptional component to oncogenesis [7]. Hence, understanding transcriptional dynamics is important for understanding disease pathogenesis including in cancer.

mRNA decay studies have traditionally been dominated by the use of transcriptional inhibitors such as actinomycin D [8, 4] or α-amanitin [3, 9] to arrest transcription. mRNA concentrations are then quantified at progressive time points after transcriptional arrest. However, these transcriptional inhibitors stress cells and themselves alter transcript localization, stability, and cellular proliferation, introducing unintended and confounding variables [10, 11]. More recent metabolic nucleotide labeling techniques such as with 5-bromouridine [12, 10], 4-thiouridine [3, 10], or 5-ethynyluridine [13] have overcome these limitations by providing a direct read-out of transcript decay. Here we exploit the recently described 5’-bromouridine IP chase (BRIC)-seq assay [12, 10], in which the nucleotides of newly synthesized transcripts are labeled with 5-bromouridine (BrU), to study transcriptomic stability in melanoma cells. Coupling BrU labeling with RNA-seq quantifies global RNA (mRNAs and long non-coding (lncRNAs)) stability over time without unduly perturbing the cellular environment [14, 12, 10].

MicroRNAs (miRNAs) are a class of single-stranded short noncoding RNA, typically 22 nucleotides long in the mature form [15]. miRNAs are well known post-transcriptional regulators in both health and disease [16, 17]. miRNAs tend to repress gene expression by binding to miRNA-specific sites UTR of the target mRNA in conjugation with the AGO protein-containing RNA-induced silencing complex (RISC) [18]. Altered regulation of miRNAs or their targets is common in cancers, with miRNA dysregulation shown to have both oncogenic and tumor suppressive roles [17, 19]. Further, miRNA targets tend to have shorter half-lives [5], so miRNAs have a direct impact on steady-state transcript levels and ultimately protein activity. Thus, a miRNA-induced change in transcript stability - for example, of oncogenes or tumor suppressor genes - has functional consequences [7, 20].

Here we explore the transcriptome stability of the stage IV amelanotic melanoma cell line A375 to better understand its role in melanoma development and progression. We also investigate RNA stability dynamics in the presence and absence of miR-211, which acts as a tumor suppressor in this cell line [21, 22]. We show that miR-211 decreases transcriptome stability. miR-211-dependent destabilization also downregulates several target genes and pathways involved in cancer progression. Using a combination of stability dynamics, gene expression changes, and *in silico* prediction, we decipher transcriptome-wide miR-211 targets in A375 cells. Focusing on one such target gene, *DUSP3*, we show that its miR-211-dependent downregulation modulates signaling pathways and the cancer phenotype *in vitro*. Overall, the study provides insights into the dynamic interplay between miR-211 and its targets and how these interactions contribute to melanoma formation.

## RESULTS AND DISCUSSION

### Transcriptomic stability in the A375 amelanotic melanoma cell line

We were motivated to study the transcriptome-wide half-life of A375 amelanotic melanoma cells to better understand the relationship between mRNA stability and tumor phenotype. To this end, we used BRIC-seq to quantify transcriptome-wide half-life [14, 12]. A375 cells were subjected to BRIC-seq at four timepoints (0, 1, 3, and 6 h) (Fig. 1A) to determine transcript decay rate or half-life. BrU labelled RNA was pulled down using an anti-BrdU antibody before being deep sequenced and processed to obtain read counts normalized to spike-in control RNA (see Methods). Normalized RNA reads were filtered for quality to obtain the decay profile (Fig. 1B), which was converted into half-lives (Fig. 1C, **Table S1**).

**Figure 1.**
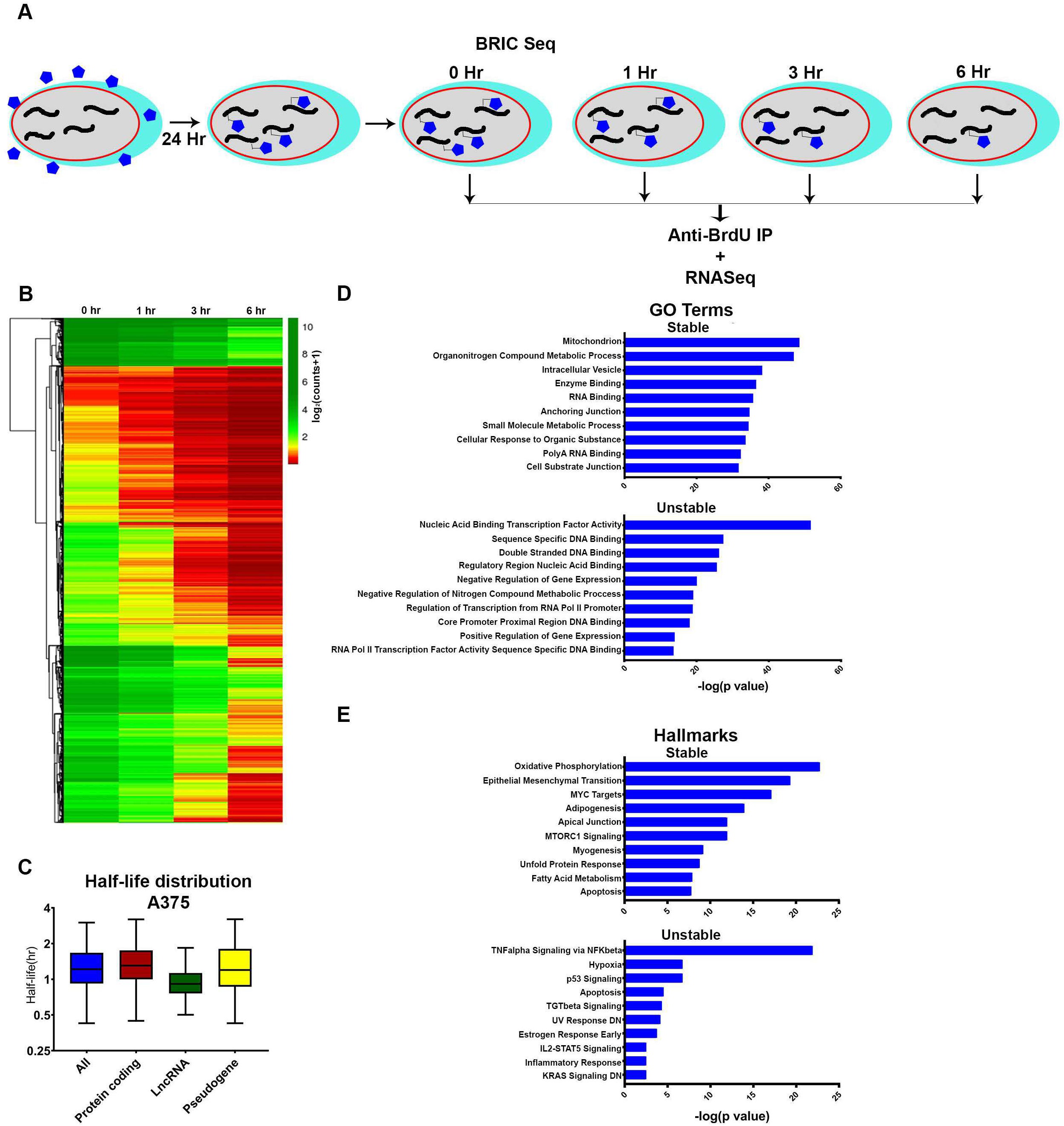
Transcriptome decay profile of the amelanotic melanoma cell line A375. A) Schematic of BRIC-seq ([10], see Methods). BrU-labeled RNA is collected at defined intervals after initial incubation to identify transcriptome-wide decay rates. B) Heatmap of normalized read counts and depicting decay profiles of all the filtered transcripts. Unsupervised hierarchical clustering represents subgroups clustered based on decay profile. C) Box plots showing the half-life distribution (in hours) of all the transcripts and transcript species as calculated from the filtered data. D) Gene ontology (GO) analysis showing top ten GO terms associated with the 5% most highly stable (top) and 5% most highly unstable (bottom) protein coding transcripts. E) Hallmark analysis showing top ten hallmarks associated with the 5% most highly stable (top) and 5% most highly unstable (bottom) protein coding transcripts.

At the transcriptome-wide level, the majority of transcripts decayed over time (Fig. 1B). Overall, transcripts had half-lives of under 2 h (mean 1.459 h, median 1.248 h) (Fig. 1C), with mRNA stability (mean 1.512 h, median 1.324 h) higher than lncRNA stability (mean 1.077 h, median 0.9364 h) but comparable to pseudogene transcripts stability (mean 1.564 h, median 1.248 h) (Fig. 1C). BRIC-seq-quantified half-lives were validated by BRIC-qPCR for select candidates, and MYC and GAPDH half-lives were within 20% of the deep sequencing estimate and half-life ratios were comparable (Fig. S1).

We next established whether transcript stability was associated with gene function in A375 cells. Gene ontology (GO) analysis of genes ranked based on half-life showed that the top 5% most stable (top 5%, high half-life) coding transcripts had a housekeeping function (e.g., mitochondrial function, metabolic processes, and enzyme binding), while the top 5% most unstable (bottom 5%, low half-life) coding transcripts tended to be important for transcriptional regulation (e.g., nucleic acid binding transcription factor activity, double-stranded DNA binding, negative regulation of gene expression; Fig. 1D), similar to observed in other genome-wide decay studies using different methodologies [23, 5, 10, 6]. With respect to hallmark signatures [24], the most stable transcripts were enriched for genes involved in epithelial to mesenchymal transition (EMT) and MYC signaling in addition to housekeeping metabolic pathways such as oxidative phosphorylation (Fig. 1E), while unstable transcripts were enriched for signaling pathways that play significant role in melanoma progression such as TNF [25, 26], p53 [27], and TGF [28, 29]. Interestingly, unstable genes were enriched for genes downregulated by UV radiation [30] and KRAS signaling [31, 32], two common drivers of melanoma progression. Overall, many pathways and processes are regulated at the RNA stability level, with the unstable decay profiles identifying highly regulated genes and pathways important in melanoma progression.

### miR-211-mediated decay targets coding and non-coding transcripts

To better understand the role of miRNAs in transcript stability and cancer progression in A375 cells, we next determined transcriptome stability dynamics in A375 cells overexpressing miR-211 (A375/211 cells), a known tumor suppressor (Fig. 2A, Fig. 2B, Fig. S2) [21, 22, 33]. miR-211 overexpression reduced the half-lives of all RNA species, especially mRNAs (mean 1.297 h, median 1.170 h, 14% reduction compared to parental cells) and pseudogenes (mean 1.365 h, median 1.187 h, ∼13% reduction compared to parental cells; Fig. 2C). While miR-211 overexpression had little effect on the half-life (Fig. 2C), transcript stability was affected in both directions but mainly towards destabilization (Fig. 2D). Our approach to look for *bona fide* miR-211 targets using multiple approaches (*in silico* prediction and RNA-seq) overcame the impact of genome-wide noise associated with the ectopic expression of miR-211.

**Figure 2.**
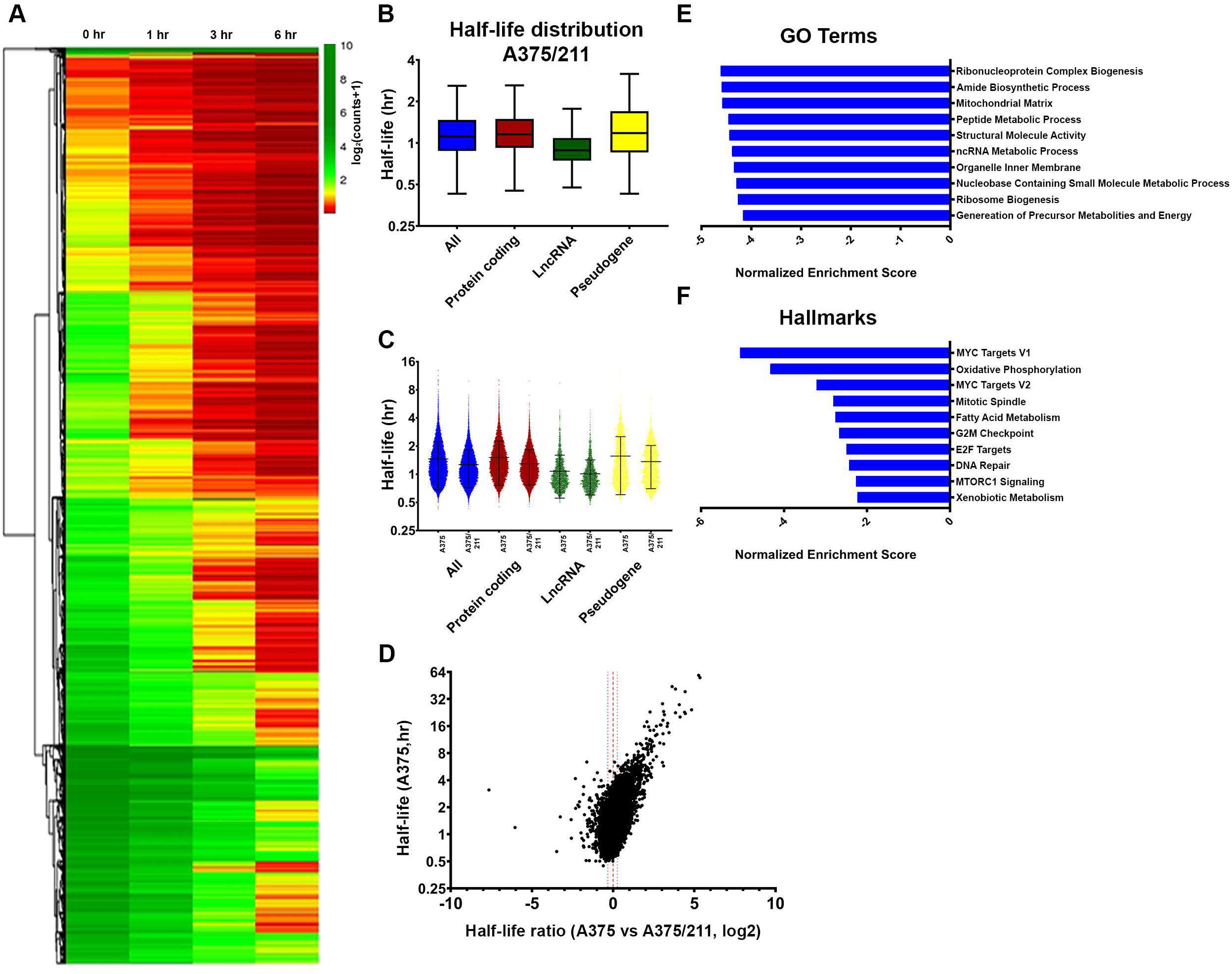
miR-211 destabilizes transcriptome stability in A375 cells. A) Heatmap of normalized read counts depicting decay profiles of all the filtered transcripts for A375/211 cells. Hierarchical clustering represents subgroups clustered based on decay profiles. B) Box plots showing the half-life distribution (in hours) of all the transcripts and transcript species as calculated from the filtered data for A375/211 cells. C) Violin plot depicting half-lives of each individual transcript for different RNA species between A375 and A375/211 cells. D) Volcano plot depicting the correlation between half-life of a transcript in control A375 cells and the change in half-life upon miR-211 induction in A375/211 cells. The majority of transcripts are destabilized upon miR-211 expression. Dashed line represent decay ratio 1 and dotted line depicts ±20% window. E) GSEA for gene ontology (GO) terms showing the top ten GO terms enriched for miR-211-driven destabilized transcripts with lower half-life (≤80%) compared to parental A375 cells. F) GSEA for hallmark terms showing the top ten hallmark terms enriched for miR-211 driven destabilized transcripts.

GO and hallmark analysis suggested that miR-211 preferentially decreased the half-lives of genes involved in known miR-211-regulated pathways (Fig. 2E, Fig. 2F) such as oxidative phosphorylation/energy metabolism [34, 22, 35, 36], lipid/fatty acid metabolism [37, 36], mitosis/cell-cycle progression/cell-cycle checkpoint [38–40], and MYC-dependent signaling [41, 42], suggesting a miR-211-dependent contribution to half-life dynamics. This altered regulation extended to the individual pathway level, with gene set enrichment analysis (GSEA) for Biocarta pathways showing that integrin signaling, involved in cell shape and mobility, was enriched in destabilized protein-coding transcripts (data not shown), in line with our previous observation of drastically altered shape of A375/211 cells compared to A375 cells [22]. Taken together, these results suggest that miR-211 drives transcript stability changes in A375 cells that are associated with functional outcomes.

### miR-211 target prediction using BRIC-seq stability data

miR-211 might destabilize transcripts through both direct (miR-211 binding to the 3’UTR of the target mRNA with the RISC complex to degrade mature transcript) or indirect (destabilized through overall changes in the regulatory state of the cell) regulation. The observed level of an individual transcript will be governed by both the degradation rate and the transcription rate.

Given that miRNAs directly degrade transcript, we reasoned that direct functionally relevant miR-211 targets will have (i) lower stability (i.e., decreased half-lives, 80%) in A375/211 cells than in parental A375 cells; (ii) lower expression levels (Fig. 3A), and (iii) miR-211 tareget sequences in their 3’UTR. Using this stringent approach, 16 protein-coding genes (Fig. 3B and Fig. 3C) had a high probability of being miR-211 targets, and indeed *PDK4* and *EFEMP2* have been shown to be direct MIR211 targets in melanoma [43, 22]. GO analysis of the selected candidates suggested that miR-211 directly targets genes related to cellular metabolism, and several of the predicted miR-211 targets have been implicated in the development and progression of other cancers; for example, *ANO5* is downregulated in thyroid cancer [44]; *GREB1* is upregulated in ovarian [45], breast [46], and prostate cancers [47]; and *EFEMP2* is upregulated in glioma [48] and colorectal cancers [49]. Some previously predicted miR-211 targets in melanoma cell lines such as *RAB22A*, *SERINC3*, *AP1S2* [50]; *NUAK1* [43]; and *BRN2/POU3F2* [51] did not meet the criteria, suggesting that the approach shows some cell type specificity. Nevertheless, combining half-life/decay data with gene expression data and existing information appears to provide a highly stringent approach for identifying miRNA targets.

**Figure 3.**
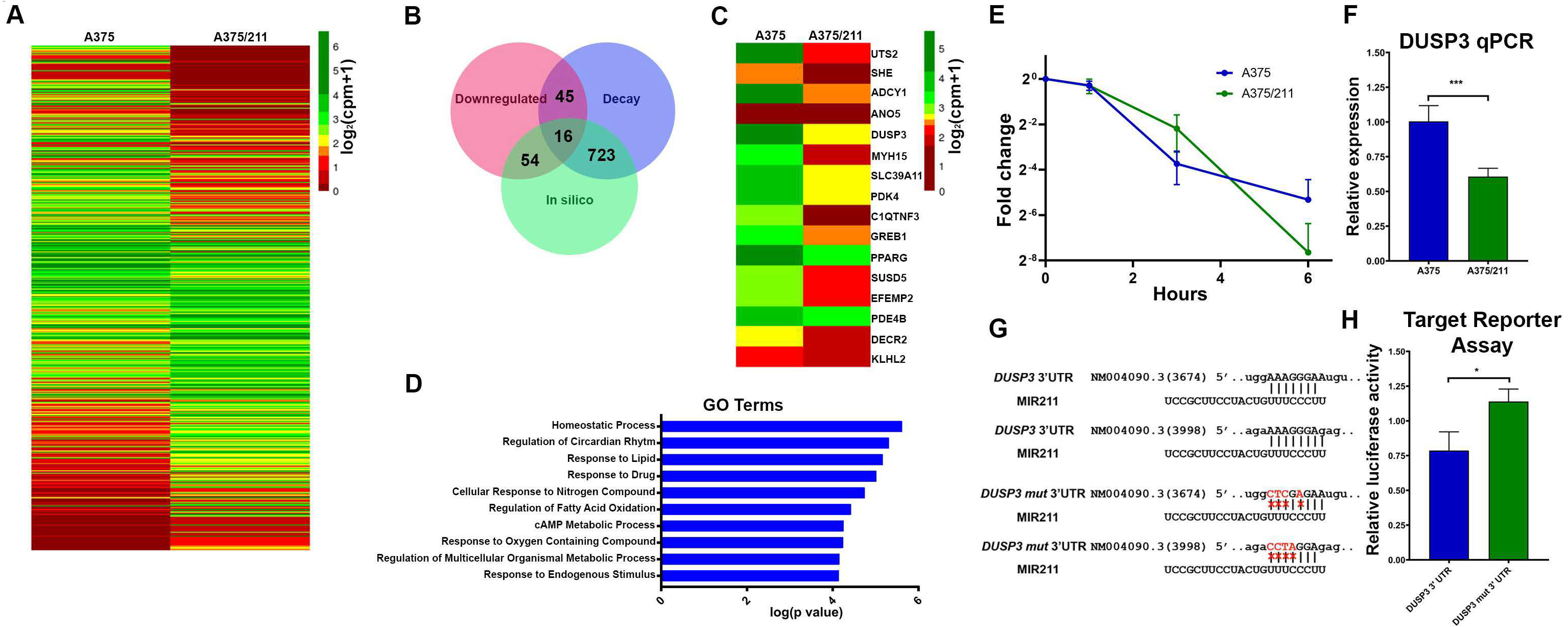
Decay dynamics helps identify miR-211 targets in A375 cells. A) Heatmap showing normalized read counts comparing expression of significantly (two-fold change, FDR <0.1) differentially expressed protein-coding genes between A375 and A375/211 cells. B) Venn diagram showing the overlap between downregulated genes, destabilized genes, and *in silico* predicted gene targets. 723 predicted targets decayed faster (half-life ratio A375 vs A375/211, >=1.2) upon miR-211 induction; 54 predicted targets were downregulated and 45 faster decaying genes were downregulated. Altogether, 16 predicted targets were destabilized and downregulated. C) Heatmap plotting normalized read counts of selected miR-211 targets based on stringent criteria of overlap between destabilization, downregulation, and *in silico* prediction. D) Gene ontology (GO) analysis showing the top ten GO terms enriched for direct miR-211 targets. E) Graph comparing decay profile of *DUSP3* between A375 and A375/211 cells as calculated via BRIC-qPCR and showing that miR-211 reduces DUSP3 transcript half-life. F) qPCR validation of *DUSP3* downregulation upon miR-211 induction in A375 cells. G) *In silico* predicted miR-211 target sites in the *DUSP3* H) *In vitro* validation of miR-211-dependent *DUSP3*3’ UTR decay via a luciferase-based mi-RNA reporter assay.

### *DUSP3* is an miR-211 target in A375 cells where it promotes cancer progression

To examine whether our approach had successfully identified a hitherto unknown miR-211 target in melanoma, we selected *DUSP3* for further validation. *DUSP3*, also known as vaccinia-H1 related (VHR) [52], has been shown to be functionally involved in various cancers including cervical carcinoma [53], prostate cancer [54], and breast cancer [55]. However, there are no reports of *DUSP3* involvement in melanoma or its regulation by miR-211.

We first validated *DUSP3* as an miR-211 target. BRIC-qPCR (t_1/2_; A375= 0.956 h, A375/211=0.775 h) (Fig. 3E) and RT-qPCR (Fig. 3F, Fig. S3) showed that miR-211 upregulated the *DUSP3* transcript decay rate and downregulated *DUSP3* expression. There were two *in silico*-predicted miR-211-binding sites in the *DUSP3*3’UTR (Fig. 3G), so we tested both sites for miR-211 targeting by transfecting cells with a luciferase vector containing native *DUSP3*3’UTR or mutated (mut) *DUSP3*3’UTR (Fig. 3G). The mutated 3’UTR abrogated miR-211-dependent downregulation of luciferase activity (Fig. 3H) compared 3’UTR.

To understand whether miR-211-mediated *DUSP3* downregulation in melanoma was functional, we next performed loss-of-function experiments with siRNA knock-down of endogenous *DUSP3* in A375 cells (Fig. 4A). While miR-211 overexpression significantly reduced proliferation compared to parental A375 cells (Fig. 4B), siRNA knockdown of *DUSP3* in A375 cells did not affect proliferation (Fig. 4B). However, *DUSP3* knockdown reduced invasion and colony formation at comparable levels to those observed in A375/211 cells compared to A375 (Fig. 4C, Fig. 4D), suggesting that *DUSP3* downregulation might contribute to miR-211-dependent regulation of cell invasion and anchorage-independent growth. *DUSP3* has previously been shown to induce cell growth and inhibit apoptosis in adenocarcinoma cells [54], and *DUSP3* depletion in HeLa cells induces cell cycle arrest and reduces cell proliferation [56]. Furthermore, *DUSP3* knockout reduces cell migration [57] and inhibits *in vitro* angiogenic sprouting [58], consistent with the oncogenic function seen here.

**Figure 4.**
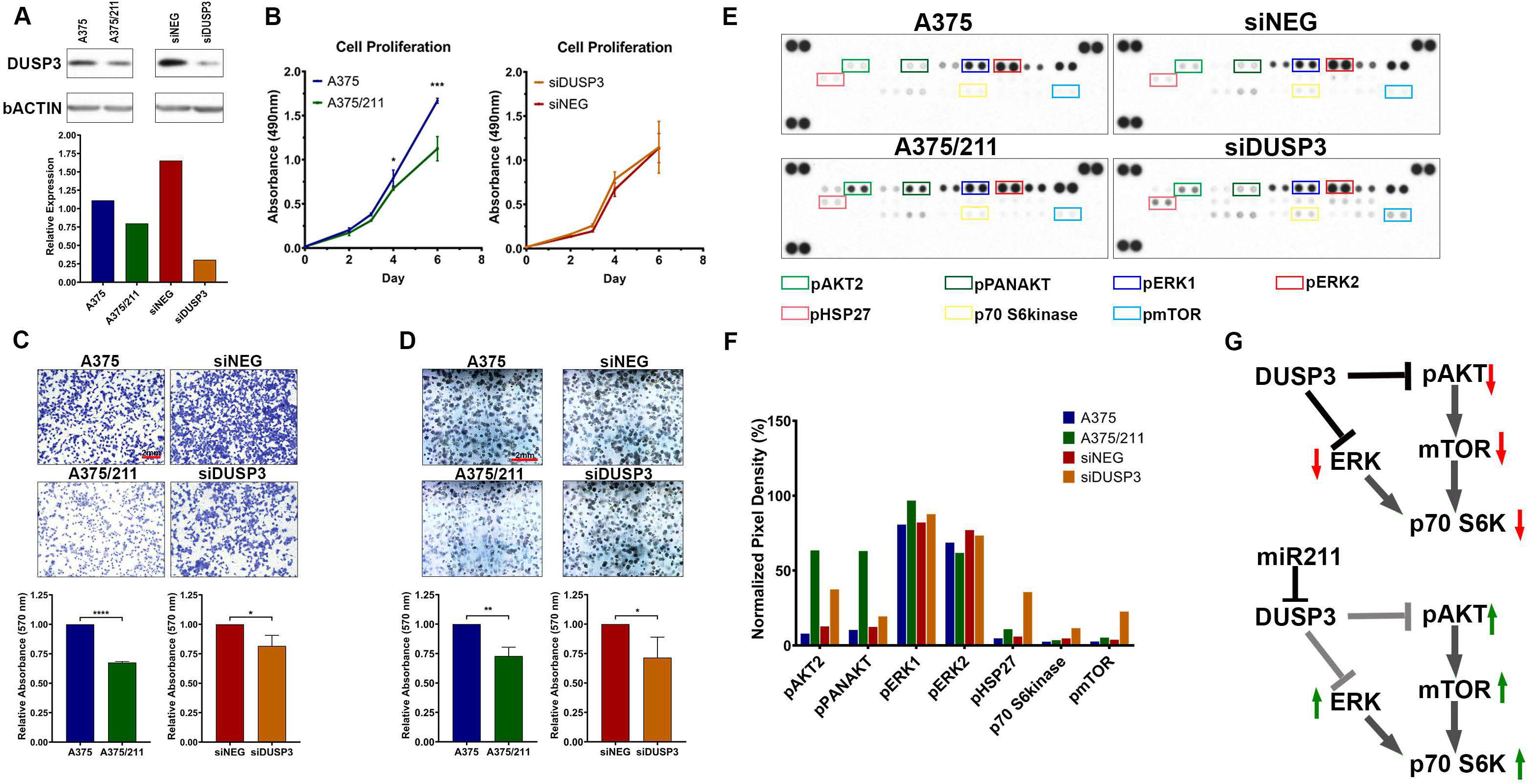
*DUSP3* downregulation reduces cell invasion and colony forming ability in A375 cells and alters MAPK/PI3K signaling. A) Western blot showing downregulation of *DUSP3* protein in A375/211 cells and validating knockdown of *DUSP3* upon transient *DUSP3* siRNA (si*DUSP3*; 48 h post-transfection) transfection in A375 background compared to control siRNA (siNEG). B) *DUSP3* knockdown does not affect A375 cell proliferation *in vitro* (right, compared to control siRNA, siNEG, based on four independent experiments). C) *DUSP3* knockdown in A375 leads reduces cell invasion (right two panels; compared to control siRNA, siNEG). Reduced cell invasion is also seen in A375/211 compared to A375 cells (left two panels). Bar graph quantifying three independent trials. D) *DUSP3* knockdown in A375 cells reduces colony formation (right two panels; compared to control siRNA, siNEG). Reduced colony formation is also seen in A375/211 compared to A375 cells (left two panels). Bar graph quantifying three independent trials. E) Human MAPK array depicting levels of various phosphorylated kinases with 8-minute exposure time. Boxed dots represent duplicate spots for identifying the indicated phosphorylated protein kinases. Bar graph representing normalized levels of selected (boxed) phosphorylated protein kinases for comparison between A375 vs. A375/211 and siNEG (control siRNA) and siDUSP3 (*DUSP3* siRNA) conditions. F) Model of *DUSP3* and miR-211-dependent dysregulation of MAPK and PI3K signaling. Significance level: * p<0.05; ** p<0.005; *** p<0.005.

*DUSP3* is a dual activity phosphatase that regulates signaling pathways by dephosphorylating target tyrosine and/or serine/threonine residues [59]. While *DUSP3* has been shown to specifically target ERK1/2 and JNK MAP kinase (MAPK) pathways in multiple cancers [58, 60, 55, 56], its role in melanoma is unknown. To investigate the specific kinase pathway(s) regulated by *DUSP3* in A375 cells, we assessed phosphorylation of various MAPK pathway members using a MAPK array upon *DUSP3* knockdown (Fig. 4E). *DUSP3* knockdown increased phosphorylation of pAKT, pERK1, pHSP27, p70 S6kinase, and pmTOR, similar to the pattern seen in A375/211 cells (Fig. 4F). While previous reports have suggested *DUSP3* targets the ERK pathway, our observation suggests that *DUSP3* also regulates PI3K/AKT signaling, not only upstream at the level of pAKT but also the downstream pathways members p70 S6kinase and pmTOR (Fig. 4G) [61]. Recent reports have suggested that HSP27 is associated with both the ERK and AKT pathways, suggesting the observed change in phosphorylation could be via these pathways [62–64]. Similar positive regulation of ERK [65] or AKT [35] by miR-211 has previously been reported in other cell lines. Taken together, these results support that miR-211 may exert some of its tumor suppressive function via downregulation of *DUSP3* and related downstream pathways to inhibit growth and invasion.

## CONCLUSIONS

Here we explored the transcriptome stability of the stage IV non-pigmented melanoma cell line A375 to better understand its role in melanoma development and progression. We also investigated RNA stability dynamics in the presence and absence of miR-211, which acts as a tumor suppressor in this cell line [21, 22]. We demonstrated that miR-211 decreases transcriptome stability and that this miR-211-dependent destabilization downregulates key target genes and pathways that leads to melanoma progression.

In general, our pathway analysis showed that genome-wide transcript stability was related to several pathways of interest in cancer and more specifically melanoma. For instance, stable transcripts were enriched for MYC targets and mTORC1 signaling, both of which are targets of the constitutively active RAS-RAF-MEK-ERK signaling pathway and important regulators of the melanoma phenotype [66–68]. Genes involved in oxidative phosphorylation and EMT also showed higher stability and, given that EMT in particular drives invasion and metastasis, implicates global transcript stability in the maintenance of critical cancer hallmarks [69–71]. Unstable genes were enriched for apoptosis and p53 signaling pathways, suggesting that global instability could disrupt critical tumor suppressive pathways.

miR-211 also had a global impact on transcript stability through direct targeting and indirect physiological changes: miR-211 directly targeted and destabilized A375 homeostatic and metabolic processes and indirectly destabilized genes involved in cell cycle, DNA repair, and MYC-dependent regulation. Interestingly, miR-211 destabilized oxidative phosphorylation in A375 cells, a previously described metabolic switch function of miR-211 in melanotic and amelanotic melanomas [36, 35, 22].

One important observation from the decay rates dynamics is the abundance of non-target-specific interactions upon miR-211 overexpression. Unfortunately, since this was a first proof of concept study using this approach, it is difficult to establish the exact cause of this observation. However, other genome-wide miRNA target identification approaches such as AGO2-RIP-seq also result in non-target-specific interactions. For instance, Meier et al. [72] investigated genome-wide translational inhibition upon miRNA (miR-155) expression and also observed indirect mRNA and miRNA expression profile alterations that contributed to the observed phenotype. While this study did not investigate transcriptome wide half-life dynamics, AGO2-RIP-seq and RNA-seq analysis suggested considerable non-target specific interactions and regulation. RNA-seq analysis of total cell lysates suggested that only 21.5% of downregulated (total 339) targets were predicted based on miR-155 binding sites alone. In addition, 11.6% of 121 upregulated genes were predicted miR-155 targets, suggesting context-dependent regulation. RIP-seq following AGO pull-down showed that 100 transcripts were enriched upon miR-155 induction and, of those, only 67 were predicted targets and only nine of those predicted targets showed regulation by decay. Similarly, Kutsche et al. [73] also found that out of 127 differentially AGO2-enriched and upregulated genes upon miR-124 deletion, only 98 were predicted targets. Taken together, these studies and out data suggest that miRNA-dependent regulation of targets without predicted binding sites could either be a by-product of miRNA overexpression that overloads AGO2 RISC with the candidate miRNA or could be a genuine interaction via currently unknown mechanisms.

Finally, we successfully exploited a genome-wide transcript half-life changes assay to identify context-dependent transcriptome-wide miRNA targets. Our combinatorial approach of using half-life ratios, differential expression, and *in silico* prediction proved to be a stringent method to identify functionally relevant and novel miRNA targets. We show that the novel direct miR211 target *DUSP3* has an oncogenic role in A375 melanoma cells. Finally, our results support a role for miR211-dependent regulation of *DUSP3* in regulating melanoma growth and progression by modulating MAPK/PI3K signaling.

## Materials and methods

### Cell culture

The human stage IV amelanotic melanoma cell line A375 (ATCC Number: CRL-1619) and the derived miR-211-overexpressing A375/211 cell line [21] were grown in Tu+ medium as previously described [21].

### BRIC-Seq and BRIC-qPCR

A375 and A375/211 cells were incubated with metabolic bromouridine (BrU) for 24 h and then used as per the BRIC kit (RN1007/1008; MBL International, Woburn, MA) protocol. Briefly, labeled medium was replaced with fresh medium and cells were collected at 0, 1, 3, and 6 h post labeling. Total RNA was extracted using the Direct-zol RNA miniprep kit (R2060, Zymo Research) mixed with external spike-in labeled RNA according to the manufacturer’s recommendation, before pulling down labeled RNA with an anti-BrdU antibody (BRIC kit). Bound RNA was eluted and followed by RT-qPCR or deep sequencing.

### RNA extraction, cDNA synthesis, and qPCR

Total RNA was isolated from cells using the Direct-zol RNA miniprep kit (R2060, Zymo Research), with subsequent quantification using Nanodrop (Thermo Fisher Scientific). TaqMan probes were used to quantify *DUSP3* (Hs01115776_m1) using *GAPD* (Hs99999905_m1) and *ACTB* (Hs1060665_g1) as control. Standard qPCR primers were used for *GAPDH*, *MYC*, and BRIC-spike in control for BRIC-qPCR. TaqMan probes were used to quantify levels of miR-211 (TM: 000514) and RNU48 (TM: 001006) expression was used as control. Fold change was obtained from the Ct value comparison. Fold change from different condition was compared using two-tailed Student’s t-test assuming equal variance.

### Northern blot analysis

Total 20µg of RNA from A375 and A375/211 cells was separated on 15% denaturing polyacrylamide Novex^TM^ TBE urea gel (Invitrogen), blotted on to BrightStar^TM^-plus positively charged nylon membrane (Invitrogen) and hybridized with UltraHyb-Oligo buffer (Invitorgen). The blots were then hybridized with DIG labelled miRCURY LNA miRNA detection probes specific for miR-211 (339111 YD00612245-BCH) and U6 (339111 YD00699002-BCH). The blots were washed with 2X SSC buffer and 0.5% sodium dodecyl sulfate (SDS) and processed for autoradiography.

### RNA-seq

RNA sequencing was performed at the Genomics Core at the Sanford Burnham Prebys Medical Discovery Institute, Orlando, FL. Before sequencing, total RNA quality was assessed with the Agilent Bioanalyzer Nano Assay (part no. 5067-1512) (Agilent Technologies). RNA-seq libraries were constructed using the Illumina TruSeq Stranded Total RNA Library preparation Gold kit (catalog no. 20020598) (Illumina Inc.) as per the instructions. The quality and quantity of the libraries were analyzed using the Agilent Bioanalyzer and Kapa Biosystems qPCR (Sigma Aldrich). Multiplexed libraries were pooled, and single-end 50 base-pair sequencing was performed on one flow-cell of an Illumina HiSeq 2500.The RNA-seq data is available on GEO, accession number GSE126784.

The reads were mapped to GRCh38/hg38 human genome assembly p12 (release 28, www.gencodegenes.org) using HISAT2 and annotated using corresponding release version gencode comprehensive annotation file. Mapped reads were quantified using StringTie to obtain FPKM values, which were converted to read counts using prepDE.py script (provided in StringTie manual). For differential gene expression analysis, read counts from two independent samples for A375 and A375/211 each were analyzed by DESeq2 script in R [74].

### Decay rate analysis

For BRIC-seq decay analysis, read counts from four time points (0, 1, 3 and 6 h) were normalized to that of spike-in RNA counts to obtain normalized read counts. Normalized read counts at time 1/3/6 h were scaled with respect to 0 h reads. The decay constant was obtained from normalized and scaled read counts by fitting it to a linear model (log(y)=x+c) using R. Data with positive decay constant or R-squared values <0.75 in both A375 and A375/211 conditions were removed from further analysis. Decay constant was converted into half-life from relation *t*_*1/2*_=*-0.693/decay_constant*.

For BRIC-qPCR decay analysis, fold changes in genes were obtained with respect spike-in RNA at each time point. Fold changes at time points 1, 3, and 6 h were then scaled to that at 0 h and fitted to an exponential curve to obtain the decay constant, which was then converted to half-life as described above.

### GSEA and Gene Ontology (GO) analysis

Genes selected for GO term and hallmark analysis were subjected to functional annotation using the GSEA web-based application (broad.mit.edu/gsea/) [24]. GSEA for decay ratio was conducted with GSEAPreranked list, with rank determined by decay rate ratio between A375 vs A375/211.

### microRNA reporter assay

The 3’UTR sequences of *DUSP3* (NM004090) mRNA genes were amplified by PCR using the primers (forward: AGTGGCAGG<u>GCGGCCGC</u>CACTGCCGGGAAGGTTATAGC, reverse: GGTGATGATG<u>ACCGGTA</u>GCGAGTCCAATTGCTTCATGTG) using A375 genomic DNA. The forward and reverse primer contained 5’UTR Notl or 3’ AgeI (underscored sequence), respectively. The PCR product was cloned into pcDNA6/luc/NP3 into a NotI-AgeI site using the Clontech In-Fusion kit (Clontech Laboratories, Mountain View, CA). To remove the two putative MIR211 sites, mutated *DUSP3*3’UTR was engineering via site-directed mutagenesis using the primers: first site (XhoI site) CACCAACATT<u>CTCGAG</u>CCAAACCATCCATCACCATG, reverse: GATGGTTTGG<u>CTCGAG</u>AATGTTGGTGCCTTTTGTGCC; second site (AvrII site), forward: TGGTAGCTCT<u>CCTAGG</u>TCTGTCCAGAAAAATTCACTC, reverse: TCTGGACAGA<u>CCTAGG</u>AGAGCTACCAAATTGTTT. The mutated insert was cloned into the same insertion site of a luciferase vector. 100,000 HEK293T cells were transfected with 33ng *DUSP3* 3’UTR or mutated 3’UTR (*DUSP3*mut) with 200 ng of miR-211 overexpression construct or empty vector along with 5 ng of pRL-CMV plasmid vector encoding Renilla luciferase gene using FuGENE transfection reagent (Promega, Madison, WI). After 48 h incubation, cells were washed in PBS and lysed using cell lysis buffer (Promega). The lysate was quantified for firefly and Renilla luciferase activity using the Dual-Glo Luciferase assay system (Promega) to determine miR-211-dependent target degradation/transcriptional inhibition. Firefly luciferase activity of each 3’UTR in each case, with and without miR-211 overexpression, was first normalized with respect to that of Renilla luciferase activity of the cell lysate. Then, firefly luciferase activity of each 3’UTR in the presence of miR-211 was normalized with that of control transfections with empty vector lacking miR-211.

### *DUSP3* siRNA knockout

A375 cells were transfected with siRNA against *DUSP3* (si*DUSP3*) mRNA (s4371, Ambion) to achieve downregulation of *DUSP3* expression. Negative control siRNA (siNEG) (AM4635, Ambion) was used as a control.

### Cell proliferation assay

Cells were plated in triplicate at 5000 cells per well into 96-well plates for the various time points indicated in the figures and legends. Cell proliferation was assessed using the CellTiter96 Aqueous One Solution Cell Proliferation Assay (MTS) kit (Promega).

### Cell invasion assay

Cell invasion assay was performed using Corning BioCoat Matrigel Invasion Chambers (Discovery Labware) according to the manufacture’s protocol. Briefly, cells were starved in FBS-free medium for 24 h before being transferred into the upper chambers in serum-free medium (50,000 cells per chamber). Medium containing 10% FBS was added to the lower chambers. Cells were incubated for 48 h at 37°C. To quantify cell invasion, migrated cells were stained with 0.5% crystal violet dye. Excess staining was removed by rinsing with water and subsequently the membrane was dried before taking images. Dye from stained cells was extracted using methanol, and optical density was measured at 570 nm on a spectrophotometer to quantify cell invasion intensity.

### Western blotting and MAPK phosphoarray

Whole cell extracts from A375 cells, A375 cells transfected with siNEG or siDUSP3, and A375/211 cells were fractionated by SDS-PAGE and transferred to a polyvinylidene difluoride membrane using a transfer apparatus according to the manufacturer’s protocols (Bio-Rad Laboratories). After incubation with blocking buffer containing 5% BSA in TBST (10 mM Tris, pH 8.0, 150 mM NaCl, 0.5% Tween 20) for 60 min, the membrane was washed once with TBST and incubated with antibodies against DUSP3 (4752, Cell Signaling Technology β-actin (4967, Cell Signaling) diluted in blocking buffer at 4°C for 16 h. Membranes were washed three times for 10 min and incubated with a 1:3000 dilution of horseradish peroxidase-conjugated anti-mouse or anti-rabbit antibodies for 2 h. Blots were washed with TBST three times and developed with the ECL system (Amersham Biosciences) according to the manufacturer’s protocols.

To determine the phosphorylation level of various MAPK pathway members, the phospho-MAPK array assay was performed using the human phospho-MAPK array kit (R&D Systems) according to the provided protocol.

## AUTHOR CONTRIBUTIONS

PJ and TS performed and analyzed BRIC-seq experiment

PJ and AB performed and analyzed IHC experiments

BL constructed *DUSP3* and DUSP3mut constructs

PJ performed and analyzed rest of the experiments PJ and RJP wrote the manuscript

## ACKNOWLEDGEMENTS

We thank John Marchica at Sanford Burnham Prebys Medical Discovery Institute Analytical Genomics core for his support with RNA sequencing. We also thank Dr. Haritha Kunhiraman, JHU, for her help with the northern blot analysis. Part of this research project analysis was conducted using computational resources at the Maryland Advanced Research Computing Center (MARCC). We would like to thank Ms. Tiffany Casey for final formatting of the manuscript.

---

**Figure S1.**
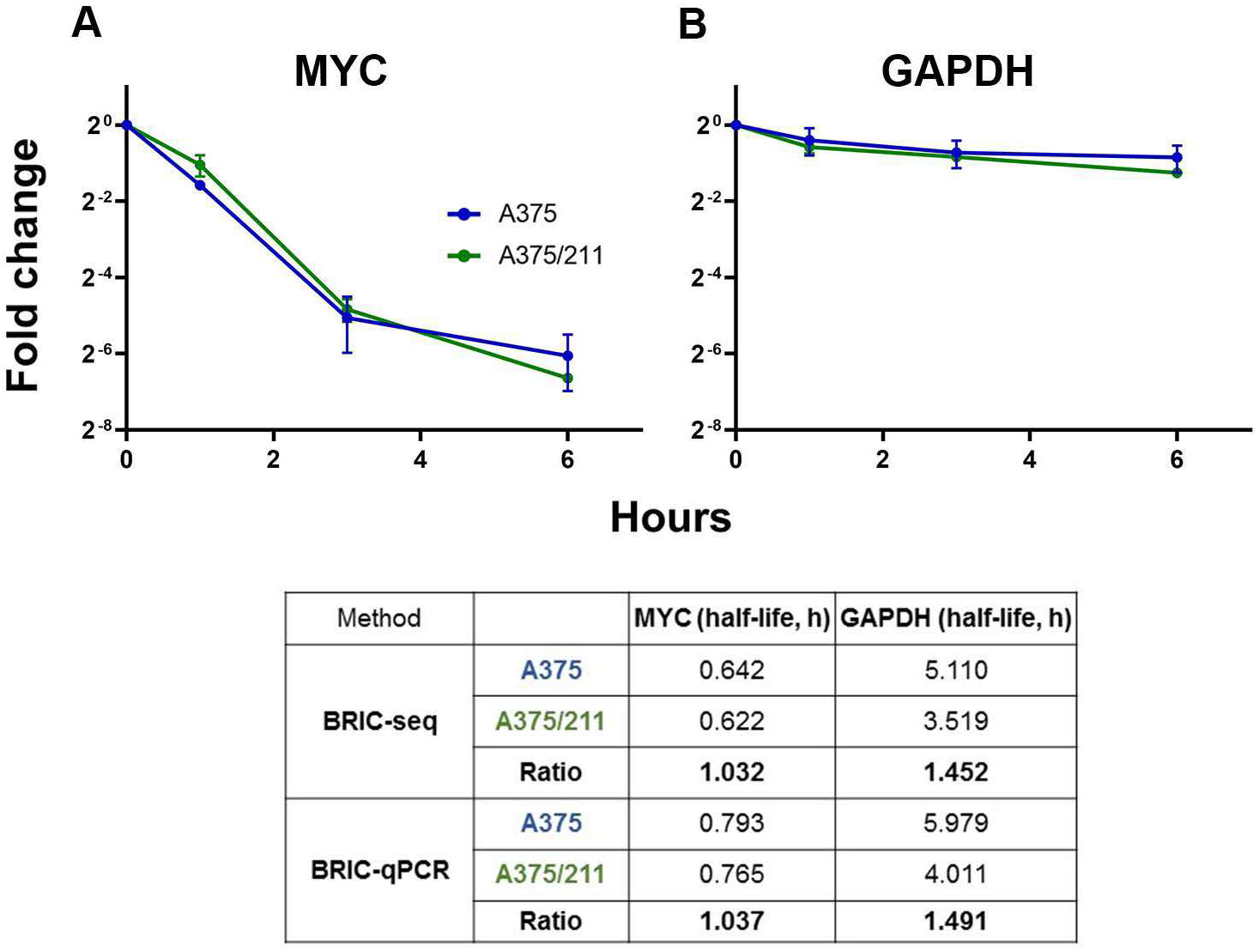
BRIC-qPCR validation of BRIC-seq data. A) *MYC* and *GAPDH* decay rates were validated by BRIC-qPCR performed at time 0, 1, 3, and 6 h. Table shows the comparison of half-lives obtained from BRIC-seq and BRIC-qPCR data for A375 and A375/211 conditions.

**Figure S2.**
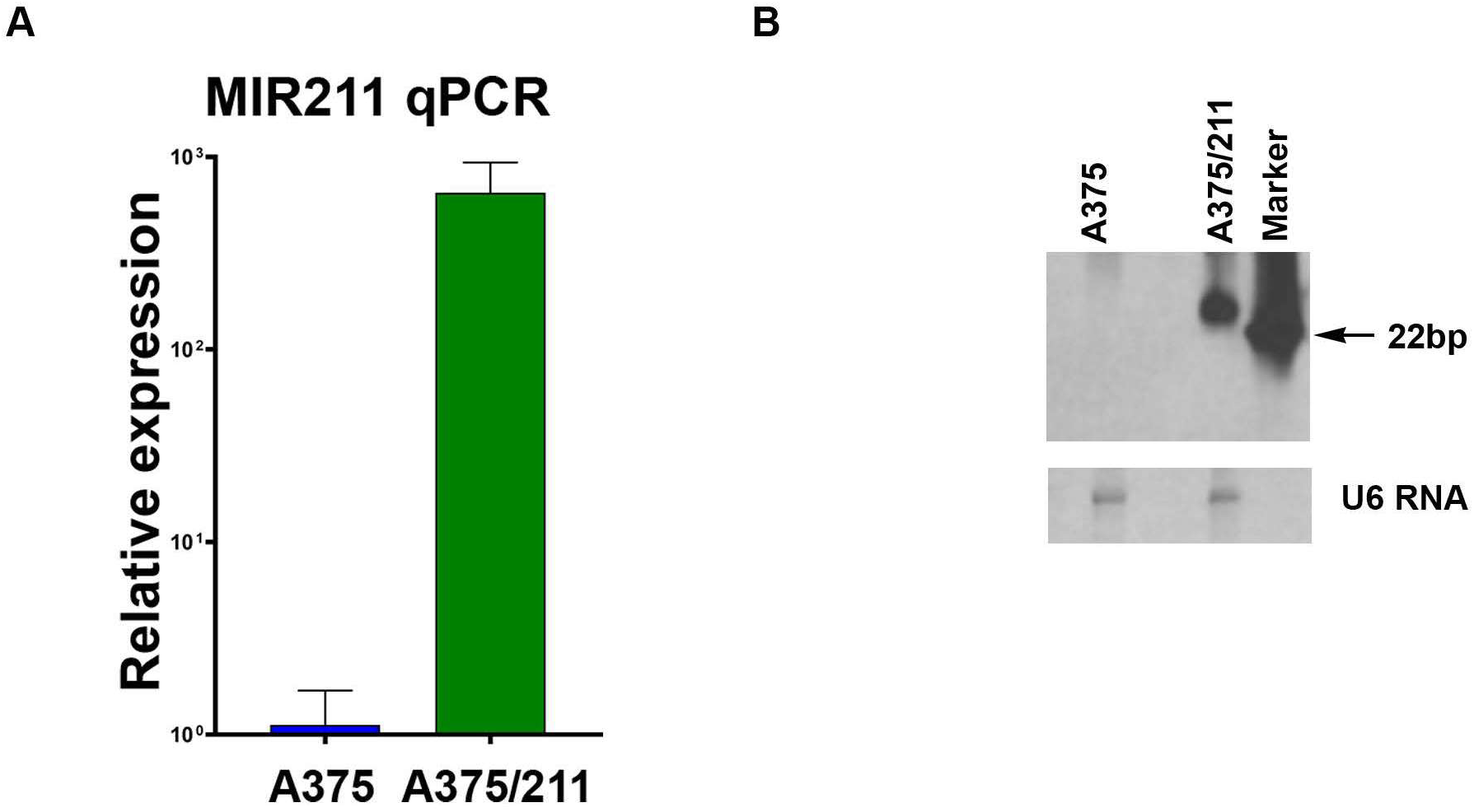
qPCR and northern blot validation of miR-211 induction in A375 cells. A) qPCR experiment showing miR-211 expression levels in A375 (average Ct: 35.1) and A375/211 (average Ct 25.6) cells. * p<0.05 B) Northern blot analysis also shows miR-211 overexpression in A375/211 cells compared to parental A375 cells. Levels of control U6 RNA were unaltered.

**Figure S3.**
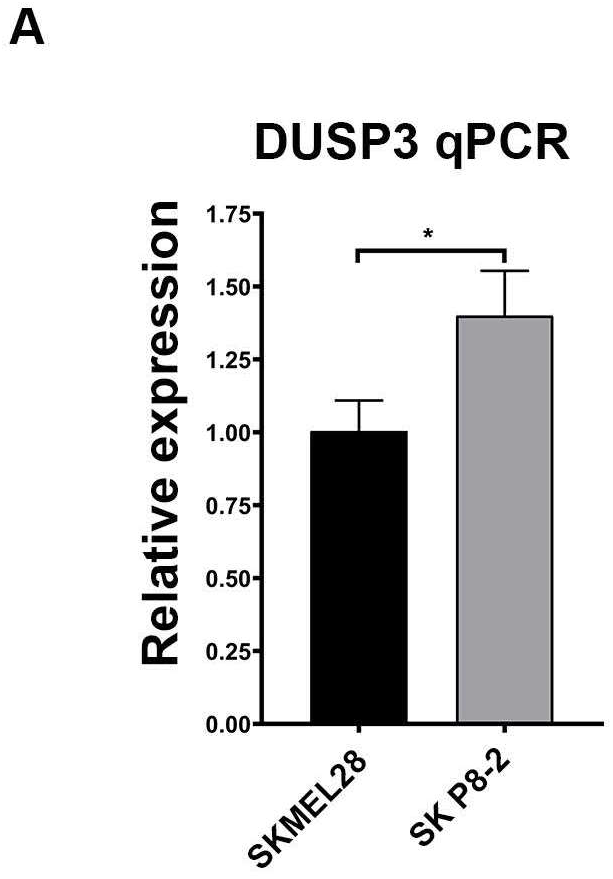
miR-211 regulates *DUSP3* in melanotic melanoma. A) qPCR validation of *DUSP3* upregulation on miR-211 deletion in SKMEL28 (SK-P8-2) compared to parental SKMEL28 cells. * p<0.05

